# Functional identification of cis-regulatory long noncoding RNAs at controlled false-discovery rates

**DOI:** 10.1101/2022.09.18.508380

**Authors:** Bhavya Dhaka, Marc Zimmerli, Daniel Hanhart, Mario Moser, Hugo Guillen-Ramirez, Sanat Mishra, Roberta Esposito, Taisia Polidori, Maro Widmer, Raquel García-Pérez, Marianna Kruithof-de Julio, Dmitri Pervouchine, Marta Melé, Panagiotis Chouvardas, Rory Johnson

## Abstract

A key attribute of some long noncoding RNAs (lncRNAs) is their ability to regulate expression of neighbouring genes in cis. However, such ‘cis-lncRNAs’ are presently defined using ad hoc criteria that, we show, are prone to false-positive predictions. The resulting lack of cis-lncRNA catalogues hinders our understanding of their extent, characteristics and mechanisms. Here, we introduce TransCistor, a framework for defining and identifying cis-lncRNAs based on enrichment of targets amongst proximal genes. TransCistor’s simple and conservative statistical models are compatible with functionally-defined target gene maps generated by existing and future technologies. Using transcriptome-wide perturbation experiments for 268 human and 134 mouse lncRNAs, we provide the first large-scale survey of cis-lncRNAs. Known cis-lncRNAs are correctly identified, including XIST, LINC00240 and UMLILO, and predictions are consistent across analysis methods, perturbation types and independent experiments. Our results indicate that cis-activity is detected in a minority of lncRNAs, primarily involving activators over repressors. Cis-lncRNAs are detected by both RNA interference and antisense oligonucleotide perturbations. Mechanistically, cis-lncRNA transcripts are observed to physically associate with their target-genes, and are weakly enriched with enhancer-elements. In summary, TransCistor establishes a quantitative foundation for cis-lncRNAs, opening a path to elucidating their molecular mechanisms and biological significance.

## INTRODUCTION

The first characterised long noncoding RNAs (lncRNAs), *H19* and *XIST*, were both found to have cis-regulatory activity: their perturbation by loss-of-function (LOF) led to increased expression of protein-coding genes encoded “in cis” - i.e. within a relatively short linear distance on the same chromosome (1, 2). Protein-coding genes whose expression responds to lncRNA LOF are considered “targets” of that lncRNA, while the direction of this change (up or down) defines the lncRNA as a “repressor” or “activator”, respectively. Since then, many more cis-regulatory lncRNAs (cis-lncRNAs) have been reported (3, 4). Conversely, other lncRNAs have no apparent positional preference for their targets and are termed ‘trans-lncRNA’ (5). This cis/trans duality provides a fundamental framework for understanding regulatory lncRNAs (6), yet the global prevalence of cis- and trans-regulatory lncRNAs remains poorly defined.

Within reported cis-lncRNAs there appears to be diversity in terms of regulatory activity (activators and repressors), distance of the target (ranging from one hundred base pairs to hundreds of kilobases) (4) (7) and number of targets (one to many) (4) (8). Two interrelated molecular mechanisms have been proposed: enhancer elements and chromatin looping (9). Some cis-activating lncRNAs, termed “enhancer lncRNAs” (e-lncRNAs), have been found to overlap DNA-encoded enhancer elements (9–12), similar to lncRNAs more generally (13). The expression and splicing of the e-lncRNA transcripts correlate with enhancer activity, implying that RNA processing somehow promotes target gene activation. Similarly, it has been proposed that cis-lncRNAs find their targets via spatial proximity, determined by chromatin looping or within the confines of local topologically-associating domains (TADs) (14). In contrast, trans-acting lncRNAs are thought to diffuse through the nucleus or cytoplasm and find their targets via molecular recognition, for example by hybridisation (15). An attractive corollary of these models is that cis-regulatory lncRNAs may act via non-sequence-dependent mechanisms, perhaps involving phase separation (16, 17) and local concentration gradients (18). It has recently been posited that lncRNAs proceed through an evolutionary trajectory commencing with fortuitous cis-regulatory activity before acquiring targeting capabilities and graduating to trans-regulation (19). Nonetheless, these conclusions are drawn from piecemeal studies of individual lncRNAs, and a holistic view of cis- and trans-lncRNAs, the features that distinguish them, and resulting clues to their molecular mechanisms and biological significance, await a comprehensive catalogue of lncRNA regulatory modes.

Regulatory lncRNA catalogues will require a rigorous and agreed definition for cis-lncRNAs, which is presently lacking. Until now, they have been defined simply by the existence of ≥1 proximal target. Targets are defined as those whose expression changes (even weakly) in response to lncRNA LOF, as measured using single-gene (RT-PCR) or whole-transcriptome (RNA-seq, CAGE, microarray) techniques (3, 5, 20). “Proximity” is defined on a case-by-case basis, using a wide range of windows spanning 10^2^ to 10^5^ bp (7). A single proximal target is usually considered sufficient. The problem with this approach is that, as the total number of targets and/or cis-window size increase, so will the chance of observing ≥1 cis-target gene by random chance. For example, consider a lncRNA having 10 proximally-encoded genes, and 2000 targets genome-wide (10% of all protein-coding genes); one would expect to observe 1 proximal target by random chance alone (10% of 100). Therefore, the conventional “naïve” definition of cis-lncRNAs, where key parameters of global target number, window size and target definition remain unconsidered or undefined, suffers from an inherent risk of false-positive predictions.

In this study, we consider cis-lncRNAs from a quantitative perspective. We show that conventional definitions are prone to high false positive rates. We introduce statistical methods for the definition of cis-lncRNAs at controlled false discovery rates and use them to classify regulatory lncRNAs across hundreds of perturbation datasets. The resulting catalogue of cis-lncRNAs enables us to evaluate hypotheses regarding their molecular mechanisms of action.

## MATERIAL AND METHODS

### TransCistor

TransCistor was developed under the R statistical software (v4.0). Gene locations were extracted from GENCODE annotation file in GTF format (v38 for humans, v25 for mouse) (21) and were converted into a matrix. The TransCistor input consists of a “regulation file”, containing all genes and a flag indicating their regulation status: 1 (upregulated after perturbation; repressed by the lncRNA), −1 (downregulated after perturbation; activated by the lncRNA) or 0 (not target). Regulation status can be defined by the user, and here is based on differential expression after lncRNA perturbation. The perturbed lncRNA itself is removed from the regulation file to avoid false positive predictions. Results are visualized with ggplot2 (v3.3.5), ggpubr (v0.4), pheatmap (v1.0.12) packages and custom in-house generated scripts.

TransCistor includes two modules; Digital and Analogue. TransCistor-digital defines cis*-*lncRNAs based on the statistical overrepresentation of proximal targets, defined as targets in the same topologically associated domain (TAD) as the lncRNA. Membership of a TAD is defined based on a gene’s TSS. Digital TransCistor utilizes a collection of TADs for human and mouse cell types accessed via the 3D-Genome Browser (22). By default, for each cell type, TransCistor identifies the lncRNA TAD and estimates the number of proximal (within TAD) and distal (outside TAD) targets / non-targets, separately for activated and repressed genes. Then, it tests for the overrepresentation of proximal targets over distal targets using the twoby2Calibrate R package. Statistical significance is estimated based on the mid-p-value calibrated Fisher’s test, for each TAD dataset/cell type. Users may use pre-calculated TAD maps employed here, or else employ TAD files of their choice in both the standalone and webserver versions of TransCistor-digital. The p-values for all the cell types are then integrated by their harmonic mean. The p-values are corrected for multiple hypothesis testing using the False Discovery Rate (FDR) method and taking into account the experiments which show at least one proximal target. The user also has the option to perform a cell type specific analysis.

TransCistor-analogue evaluates whether the mean distance of targets from the same chromosome is closer than random chance. Distance is defined by TSS to TSS. Analysis is performed separately for activated and repressed targets. Then, the random distribution is calculated, by randomly shuffling the regulation flags on genes within the same chromosome and recalculating the test statistic each time. By default, 10,000 simulations are performed. Finally, the empirical p-value is calculated from the proportion of simulations with a test statistic less than the true value.

Both modules of TransCistor are available as a standalone R package along with all regulation files (https://github.com/gold-lab/TransCistor) and Rshiny webserver (https://transcistor.unibe.ch/). The input comprises metadata about the lncRNA, and a regulation file containing target gene information that can be readily derived from any transcriptome-wide data including RNA-sequencing, CAGE and microarray experiments.

### Collecting and processing perturbation datasets

The FANTOM perturbation datasets were downloaded from the Core FANTOM6 repository (20, 23). The differential expression results were transformed into regulation files by applying an adjusted p-value threshold of 0.05 and using custom bash scripts. The respective metadata were also downloaded from FANTOM6 and were integrated into the GENCODE annotation matrix. 31 perturbation experiments were removed because they target protein-coding genes, and an additional 19 were removed because target lncRNAs had no ENSEMBL identifier. The LncRNA2Target datasets were downloaded from the webserver (Version 2.0) (24) and targets were defined by using an adjusted p-value cutoff of 0.05. The lncRNA locations were manually obtained from the website or original publications, when necessary. The rest of the datasets were accessed through the original publications and post-processed to generate the regulation files. All regulation files are available at the project Github repository, linked above.

### TransCistor concordance score

To evaluate the consistency of lncRNA classification for TransCistor, we calculated a concordance score based on lncRNAs with more than one perturbation experiment for both modes separately. We then randomly shuffled the classification labels (cis-activator, cis-repressor, not significant, or no target/TAD found.) 1000 times to create a null distribution and recalculated the score. The actual scores for TransCistor-digital and analogue were then compared with the null distribution to assess whether they provided consistent classifications.

### Analysis of subcellular localisation and expression

The data used in the initial subcellular localization analysis (Supplementary Figure S4) was downloaded from lncATLAS (25). The list of lncRNAs considered cis- or non-cis-acting respectively is based on the predicted activity as reported in Supplementary File 1. Excluding genes with no associated ENSEMBL ID, mouse genes, and genes for which either the cytosolic or nuclear expression level was missing in the lncATLAS data resulted in a reduction from 33 to 20 cis- and 290 to 133 non-cis-acting-lncRNAs for the purpose of this comparison. The Wilcoxon rank sum test was then used to check for significant differences between cis- and non-cis-acting-lncRNAs (1-sided, alternative: cis > non-cis) for a) log-transformed cytosolic/nuclear ratios, and b) total log2 FPKM cell expression. A similar analysis with datasets of total RNA (from two cell lines) was also performed.

### LncRNA evolutionary conservation analysis

Data were obtained from the LnCompare (26) data tables. PhastCons scores were utilized for the hg38 human genome assembly to obtain conservation scores of both promoter and exon regions. Three models were employed, namely 7, 20, and 100 species.

For each category, we first classified the lncRNAs into two groups: cis and non-cis-acting (same groups as for subcellular localisation). Then, we compared the PhastCons scores between these two groups using a Welch test, with a one-sided hypothesis that cis lncRNAs have greater conservation compared to non-cis-acting-lncRNAs. The comparisons were performed separately for promoter and exon regions of the lncRNAs in each of the three models.

### Target gene expression changes

For each lncRNA, we defined the proximal and distal targets based on the considered cis-regulatory region. For Trans-Digital, we employed TAD overlap, while for Trans-Analogue, we considered the entire chromosome. In cases the lncRNA was found by both methods, we used chromosome as the reference for cis-regulation. For cis-activators, we included only the downregulated targets (−1), for cis-repressors, only the upregulated ones (1), and for non-cis-acting, we incorporated both (−1,1) values. Subsequently, we compared the absolute log2 fold change values of these proximal and distal targets upon lncRNA knockdown using the Wilcoxon rank sum test (1-sided, alternative: proximal>distal).

### Analysis of chromatin states

Chromatin states annotations were retrieved from three sources: EpiMap (27), genoSTAN (28), and dbSUPER (29) EpiMap consist of 18 chromatin states across 833 samples, genoSTAN identifies promoter and enhancer regions genome-wide across 127 samples, and dbSUPER aggregates 82234 human superenhancers from 102 cell types/tissues. The annotations were relabelled as follows: Superenhancer – dbSUPER’s superenhancers; Enhancer(1) – genoSTAN’s enhancers; Enhancer(2.1) – EpiMap’s Genic enhancer 1; Enhancer(2.2) – EpiMap’s Active enhancer 1; Enhancer(2.3) – EpiMap’s Weak enhancer; Promoter(1) – genoSTAN’s promoters; and Promoter(2) – EpiMap’s Active TSS.

The human TSS annotations were intersected with chromatin states at several genomic windows (1 bp, 100 bp, 1000 bp, 10000 bp) and a given state-TSS intersection was counted only if it was present in more samples than a given threshold (0, 1, 5, or 10 samples). For each pair of genomic windows and filter, a contingency matrix was computed for each pair of predicted labels (*cis-*activator vs. cis-repressor, cis-activator vs. non-cis-acting, and cis-repressor vs. non-cis-acting) or the grouped label (*cis* vs. non-cis-acting), counting the number of TSSs falling into each category. Fisher’s exact test was used to compute the p-value of each contingency matrix.

### Chromatin looping analysis

HiC interaction data were obtained using the Python package “hic-straw” (v1.2.1) (https://github.com/aidenlab/straw), using human HiC datasets from Aiden laboratory (30). The binning resolution was set to 25 kb, and interaction scores were normalized by Knight-Ruiz matrix balancing method. Due to gaps in the HiC matrices, ∼7% of lncRNA: (non-/)target interactions were approximated by using a “next best” pair of bins, for which an interaction score was available, instead of the correct binning. In 6.8% of cases, this only required replacing either one of the ideal bins with a direct neighbour and for the remaining 0.2% either shifting both genes by one bin or one of the genes by two bins. An estimate for the expected interaction at a given distance was then calculated by fitting a regression model to the HiC data with the interaction score as the response and the TSS distance between the two genes as the explanatory variable. After visual inspection of the data, an asymptotic regression model was chosen for this step (‘SSasymp’ and ‘nls’ of the R base package ‘stats’ v4.0.3). Due to model limitations as well as unclear comparability of TAD-based and TAD-independent cis-regulation, only cis-lncRNAs identified by TransCistor-digital were included in this analysis. For 2/12 lncRNAs from this subgroup (RAD51-AS1, NARF-AS2), model generation failed for one or more of the cell types considered. Modelling the interaction as a function of the inverse square distance was also considered (‘glm’ also from stats). This model had the advantage of not failing for either combination of cis-lncRNA and cell type, but fit the data visually less well and it had a clear bias to underestimate the interaction in close 2D proximity and overestimate interaction further away (Supplementary Figure S7). The significance of interaction on the targeting status was then assessed by fitting a logistic regression model to predict whether a gene is a target of a given lncRNA based on the difference between observed and expected interaction (glm function also from ‘stats’).

### Biochemical interaction analysis

The All-toAll RNA-DNA interaction, were sourced from the RNA-Chrom database (https://rnachrom2.bioinf.fbb.msu.ru/experiments) (31) across various human cell models (HFFc6, HEK293T, HUVEC, MDA-MB-231, fibroblasts, and K562).

The dataset consolidates siginicficant interactions between the transcribed RNA and genomic DNA region from multiple techniques. To further streamline the dataset for downstream analysis, all cell line data was unified, retaining only unique RNA-DNA interaction coordinates. Further, to delineate specific interaction types, DNA coordinates from this dataset were intersected with both gene promoters (+-1000 TSS) and gene body regions, enabling distinct analyses of RNA-to-gene-promoter and RNA-to-gene-body interactions.

Subsequently, within the TransCistor dataset focusing on lncRNA knockdown in human cell models, a statistical analysis was conducted to asses the biochemical/physical interactions among genes classified as targets (1, −1) and non-targets (0) on the same chromosome as the identified lncRNA from the lncRNA’s Loss of Function (LOF) file (described previously).

This analysis generated a contingency matrix, comprehensively evaluating biochemical interactions among targets and non-targets across all lncRNAs, both collectively and at each individual lncRNA level. Following this, a one-sided Fisher exact test was performed to evaluate the if there was an enrichment of biochemical interactions among targets compared to non-targets within the context of these lncRNAs The Fisher exact test was performed separately for interactions occurring solely within gene promoters and for interactions spanning entire gene regions.

## RESULTS

### A quantitative, functional definition of cis-lncRNAs

To better understand the lncRNA target genes, we explored changes in the transcriptome arising from lncRNA perturbations across multiple studies. We employed a functional definition of “targets”, as genes whose steady-state levels significantly change in response to a given lncRNA’s LOF (Figure 1A). We further define targets as activated or repressed, where they decrease / increase in response to lncRNA loss-of-function (LOF), respectively. Overall we collected 488 lncRNA LOF experiments targeting 268 human lncRNAs from a mixture of sources, including the recently published datasets of ASO knockdowns in human dermal fibroblasts and induced pluripotent stem cells (iPSCs) from the FANTOM consortium (20, 23). To this we added 140 experiments for 134 lncRNAs from mouse (Figure 1B). Among these we included 6 hand-curated previously-reported cis-acting lncRNAs (*UMLILO*, *XIST* [x2 independent experiments], *Chaserr*, *Paupar* and *Dali*). 130 lncRNAs were represented by two or more independent experiments (Supplementary Figure S1A), and the median number of target genes identified per experiment was 55 (Supplementary Figure S1B).

**Figure 1:**
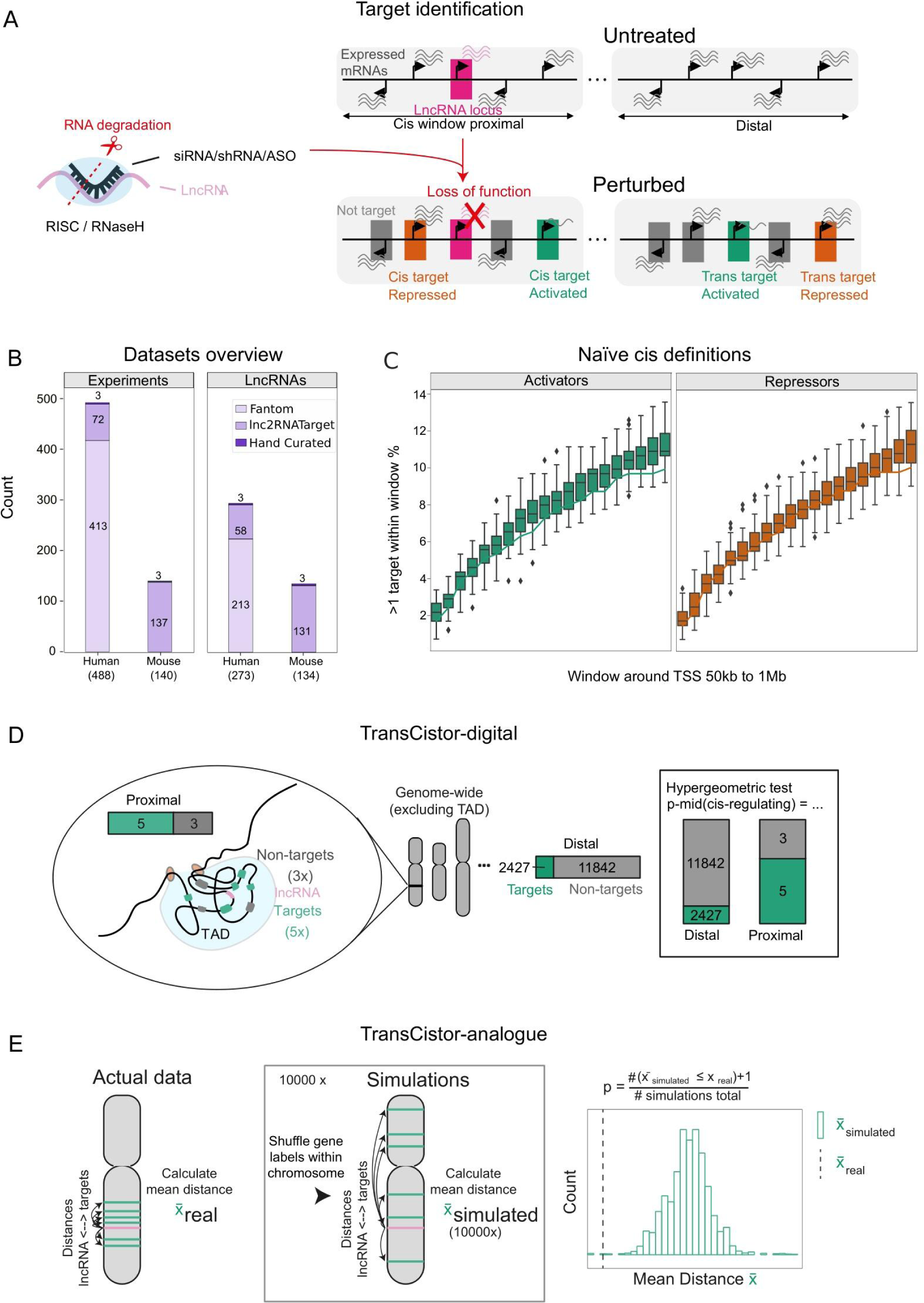
TransCistor is a quantitative framework for classifying cis-regulatory lncRNAs. A) Definition of target genes: A target gene is defined as one whose expression significantly changes after loss-of-function perturbation of a given lncRNA (pink). The direction of that change (down/up) defines the target as activated/repressed (green, orange), respectively. B) The perturbation datasets used here: Data were mainly obtained from two sources, FANTOM^47^ and Lnc2Target^48^ (x-axis). The y-axis displays the number of individual experiments (left panel) or individual lncRNA genes (right panel) (note that the difference arises from the fact that many lncRNA genes are represented by >1 experiment). Some lncRNAs are present in both datasets. C) Evaluating accuracy of naïve cis-lncRNA definition: The plot displays the number of lncRNAs classified as “cis-regulatory” using a definition of ≥1 proximal target genes (y-axis), while varying the size of the genomic window (centred on the lncRNA TSS) within which a target is defined as “proximal” (x-axis). Line: real data calculated with lncRNAs from Panel A; Boxplot: Simulations created by 50 random shuffles of the target labels across all annotated genes. D) TransCistor-digital method: TransCistor-digital evaluates the enrichment of targets (green) in proximal regions, defined as those residing within the same topologically associating domain (TAD) as the lncRNA TSS (pink) (left panel), compared to the background target rate in the rest of the genome (“Distal”) (centre panel). Cis-lncRNAs are defined as those having a significantly higher proximal target rate, defined using p-mid adjusted hypergeometric test (right panel). E) TransCistor-analogue method: A distance statistic is defined as the mean genomic distance (bp) of all targets (green) on the same chromosome as the lncRNA (pink) (left panel). 10000 simulations are performed where target labels are shuffled across genes within the same chromosome (centre panel). Cis-lncRNAs are defined as those whose real statistic (dashed line) falls below the majority of simulations (right panel).

We first evaluated the performance of the widely-used naïve definition for cis-lncRNAs, defined as ≥1 target within an arbitrary distance window. Using a range of window sizes from 50 kb to 1 Mb centred on the lncRNAs’ TSSs, we evaluated the fraction of lncRNAs that would be defined as cis-acting under this definition. This approach defines ∼2 to 12% of lncRNAs as cis-regulators (Figure 1C, line). To test whether this rate is greater than random chance, we shuffled the target/non-target labels of all genes and repeated this analysis. Surprisingly, the rate of cis-lncRNA predictions in this random data overlapped the true rates in all windows (Figure 1C, boxplots). In other words, the naïve definition of cis-lncRNAs yields high rates of false-positive predictions.

To overcome this issue, we adopted a new definition of cis-lncRNAs: *cis-lncRNAs are those whose targets are significantly enriched amongst proximal genes*. This definition has the advantage of being quantitative and statistically testable. LncRNAs that do not fulfil this criterion may be trans-acting; however, they may alternatively be genuinely cis-acting, yet obscured by large numbers of changing genes, due to technical off-targets of the perturbation method, or secondary downstream transcriptional changes. Therefore, we here term all such lncRNAs as ‘non-cis-lncRNAs’.

### TransCistor: digital and analogue identification of cis-lncRNAs

We incorporate this definition into two alternative methods for identifying cis-lncRNAs, which differ in their approach for defining cis-enrichment of targets. The first method considers proximal genes to be those whose TSS falls within a defined window around the lncRNA TSS. We developed a pipeline, TransCistor-digital, which takes as input a processed whole-transcriptome list of target genes (“regulation file”), and tests for statistical enrichment in proximal genes (Figure 1D)(Materials and Methods). Although in principle any sized window may be used, we reasoned that the most biologically meaningful would be the local TAD, in line with previous studies (32). Chromatin folding and TADs vary to an extent between cell types. Therefore, TransCistor-digital calculates enrichment across a set of experimentally-defined cell-type-specific TADs (45 human, 3 mouse) (22) and aggregates the resulting P-values by their harmonic mean.

The above TAD-window approach is effective, yet has drawbacks. Several reported cis-lncRNAs have individual targets that are not immediately adjacent (7), and might be overlooked by the digital approach. Furthermore, many lncRNAs may have no neighbouring genes in their local TAD, or no identified local TAD. Therefore, we developed an alternative method that dispenses with fixed windows, while still testing for proximal enrichment of targets. This method, TransCistor-analogue, defines a distance statistic as the mean TSS-to-TSS distance of all same-chromosome targets of a given lncRNA. Statistical significance can be estimated empirically, by generating a null distribution based on randomisation of target labels (Figure 1E). Now, cis-lncRNAs are defined as those having a distance statistic that is lower than the majority of randomisations.

We sought to test the performance of TransCistor-digital and evaluate the global landscape of cis-lncRNAs. After filtering out unusable datasets (having no targets, or no overlapping TAD), 195 datasets remained (Supplementary Table S1). We discovered 23 cis-acting lncRNAs (14 activators, 9 repressors), at a false discovery rate (FDR) threshold of 0.25 (Figure 2A-C). The majority of p-values produced by this analysis follow the null distribution, underlining the conservative statistical behaviour of TransCistor (Figure 2A,B). All cis-lncRNAs have a unique activator or repressor assignment, with the exception of *Evx1os*, which is classified as having both activating and repressing characteristics (Figure 2C). Amongst the top-ranked cis-lncRNAs is *UMLILO*, previously reported to activate multiple genes in its local genomic neighbourhood (8). *UMLILO* exhibits a significant enrichment of activated targets amongst proximal genes, which is not observed for repressed targets (Figure 2D,E).

**Figure 2:**
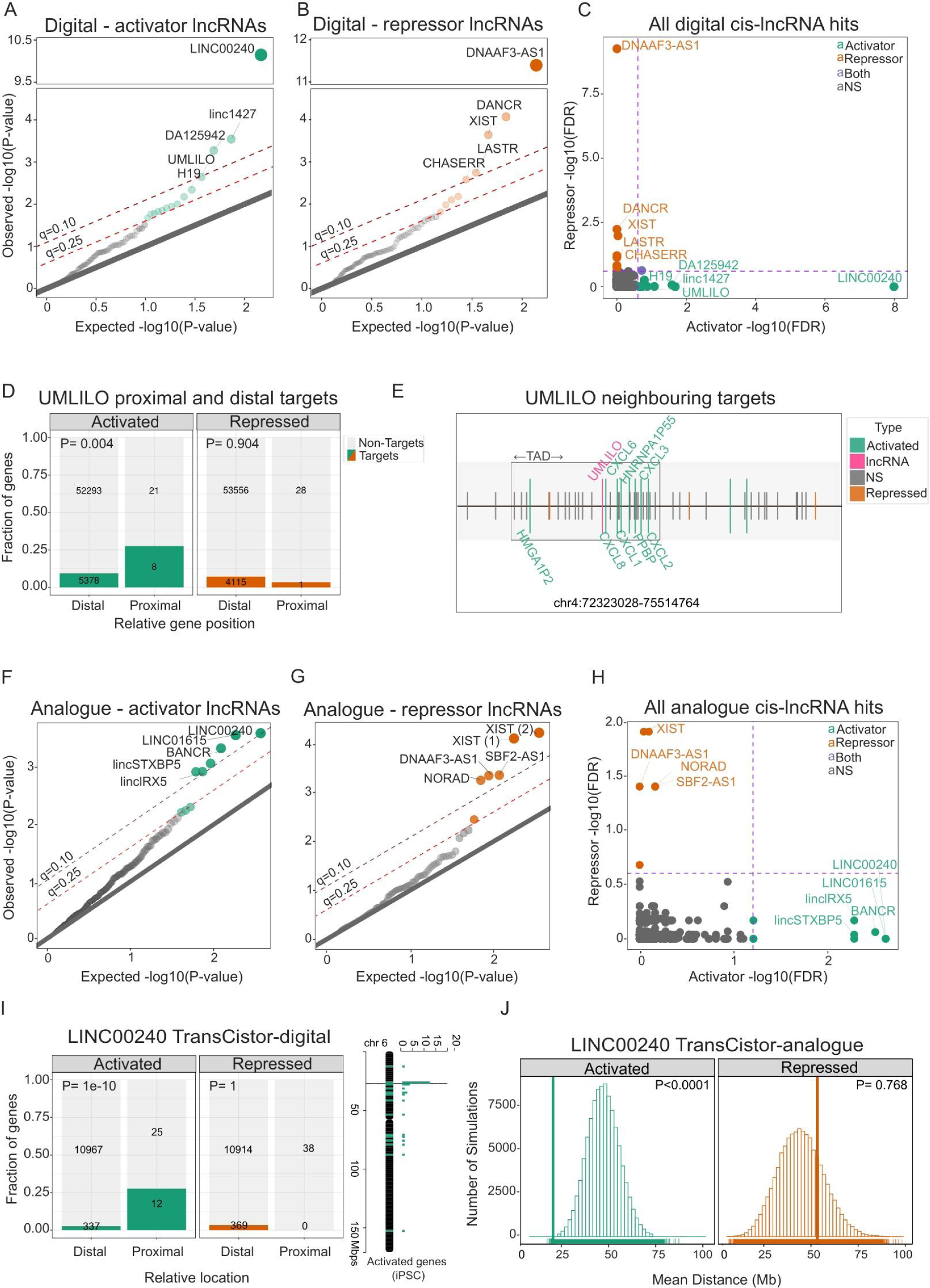
Large scale classification of cis-lncRNAs in human and mouse. A) Quantile-quantile plot displays the random expected (x-axis) and observed (y-axis) p-values for lncRNAs (points) tested for activated targets by TransCistor-digital. The grey diagonal y=x line indicates the expectation if no hits were present. Light and dark red dotted lines indicate an FDR cutoff of 0.25 and 0.10 respectively. B) As for (A), for TransCistor-digital and repressed targets. C) Comparison of activator and repressor activity detected by TransCistor-digital. For each lncRNA (points), their false-discovery rate (FDR)-adjusted significance is plotted on the x-axis (activator) and y-axis (repressor). Note the absence of lncRNAs that are both activators and repressors. D) UMLILO, an example cis-activator: The plot shows the number of genes, divided by targets / non-targets (colour / grey), location (distal/proximal) and regulation direction (activated/repressed). UMLILO is classified as a cis-activator, due to the significant excess (8) of proximal activated targets. Statistical significance (uncorrected) is displayed above. E) UMLILO genomic locus: Vertical bars denote gene TSS. Grey: non-targets; green: activated targets; pink: UMLILO. Black box: Topologically associated domain. F) As for (A), for TransCistor-analogue and activated targets. G) As for (B), for TransCistor-analogue and repressed targets. Two experiments supporting the XIST lncRNA appear in the most significant classifications represented by (1) & (2). H) As for (C), for TransCistor-analogue. I) and J) LINC00240, an example cis-activator identified by both TransCistor-digital and TransCistor-analogue. I) as for D). J) Shown is the target distance statistic (x-axis) for real data (vertical bar) and simulations (boxes). The number of simulations in each distance bin is displayed on the y-axis.

Analysis of the entire perturbation dataset by TransCistor-analogue, on the other hand, identified 15 cis-lncRNAs (9 activators, 6 repressors, FDR≤0.25) *(Supplementary Table S1)*. Statistical behaviour is good (Figure 2F,G), while cis-lncRNAs are cleanly split between activators and repressors (Figure 2H). *LINC00240* was identified as the most significant activator cis-lncRNA by both TransCistor-digital and analogue (Figure 2I,J). Similar to previous reports, we observed a strong cis-regulation of nearby histone genes (Figure 2I, Supplementary Figure S2A). (20, 23). Therefore, TransCistor correctly identifies previously-reported cis-lncRNAs.

The usefulness of these methods is further supported by their internal and external consistency. Together, the TransCistor approaches correctly identify previously-described cis*-*activators *H19* (*33*), RP11-398K22.12 (20, 23), *JPX* (*34*)*, Evx1os* (*35*), LINC01615 (36) and *DA125942* (*37*) amongst the top-ranked cis*-*activators, while both independent experiments for *XIST* are amongst the top repressors (38).

TransCistor predicted cis*-*regulatory activity for several known lncRNAs that have never been described as such in prior literature. These include *CAT2, NARF-AS2, BANCR, CD27-AS1* (*cis-*activators), and *SBF-AS1*, *LASTR, NORAD*, and *DANCR* (*cis-*repressors). However, the latter two are only identified in one out of multiple independent perturbation experiments (6 and 5 for *NORAD* and *DANCR* respectively). In the case of *DANCR*, the cis definition arises from the repression of two same-strand small RNAs (has-mir-4449, SNORA26). It is not yet clear if these results reflect false-positive or false-negative predictions. To investigate this, we merged all hits across experiments and repeated the analysis, but here we found no signal, suggesting that they are false-positive predictions. On the other hand, analysis of an independent dataset for *SBF-AS1* from different cells (A549 lung adenocarcinoma) and perturbation (siRNA), which was not included in our original dataset, yielded concordant cis-repressor prediction from TransCistor-analogue (Supplementary Figure S2B-D). Both human and mouse orthologues of *CHASERR* (ENSG00000272888) are identified as cis*-repressors* (*4*) (Figue2B). Furthermore, Chaserr is concordantly classified for independent LOF methods, knockdown by ASO and knockout by deletion (Supplementary Figure S2G). Further examples were found where ≥2 independent experimental perturbations yielded consistent predictions (*XIST* & *DNAAF3-AS1* classified as cis*-*repressors based on two separate experiments each) (Supplementary Figure S2E-F). To further test the value of TransCistor predictions, we evaluated the consistency of cis predictions between independent perturbation experiments of the same lncRNA. Using simulations to evaluate significance, we observed a significant (P<0.001) concordance between experiments for TransCistor-digital and analogue methods (Figure 3C).

**Figure 3:**
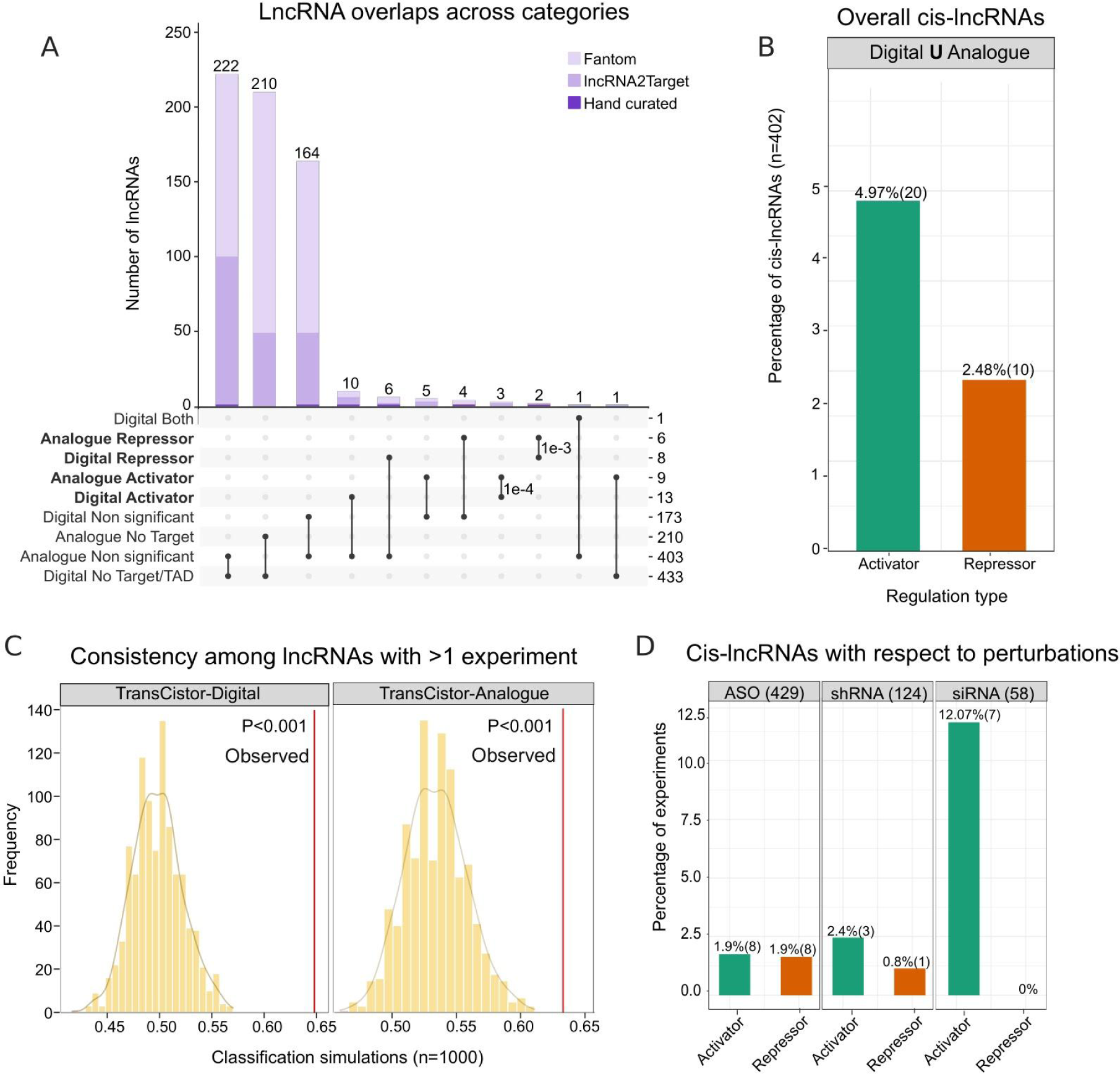
Rate and consistency of cis-lncRNA inference. A) Summary of TransCistor results across datasets. Significance of overlaps is calculated using the hypergeometric distribution. B) The rate of lncRNA genes defined to be cis-regulatory based on our analysis (union of digital and analogue). Note that one single experiment is sufficient to label a lncRNA gene as cis-regulatory. C) The concordance score is calculated for TransCistor-Digital and Analogue, based on the consistency of classification for lncRNAs with multiple supporting experiments (>1). To assess the significance, the score is recalculated for 1000 simulations with shuffled classification labels. The real concordance score, indicated by the red line, exceeds the distribution obtained from the simulated data. D) The rate of experiments defined as cis-regulatory, broken down by perturbation method.

Conversely, TransCistor failed to find evidence supporting previously reported cis*-*lncRNAs, namely *Paupar* (*39*) and *Dali* (40). Inspection of the originating microarray data revealed that, for neither case, the reported cis-target genes pass cutoffs of differential expression (Supplementary Figure S3A-D), suggesting that these lncRNAs are not cis-regulatory.

A summary of the entire set of TransCistor predictions is found in Figure 3A and Supplementary Table S2. We observed a significant overlap between the two TransCistor methods for classified activators (P=0.0001) and repressors (P=0.001), with 5 cis-lncRNAs in common (LINC00240, linc1427, DA125942 DNAAF3-AS1 and XIST). Overall, if we consider lncRNAs where at least one method in one dataset is called cis-acting, then our data implicate 7.46% (30/402) of lncRNAs as cis-regulators. When broken down by direction of regulation, we find that 4.97% (20/402) are activators and 2.48% (10/402) are repressors, with one being identified as both (Figure 3B). Henceforth, we defined the remaining 372 tested lncRNAs provisionally as ‘non-cis-lncRNAs’.

In summary, this provides a resource of cis-lncRNAs, together with multiple lines of evidence supporting the ability of TransCistor to identify true cis*-*lncRNAs.

### TransCistor cis-lncRNA identification across perturbation technologies

Our transcriptomic dataset contains a mixture of RNA interference (RNAi) and antisense oligonucleotide (ASO) LOF perturbations. While early experiments were performed using the two RNAi approaches of siRNA and shRNA, it is widely thought that these principally degrade targets located in the cytoplasm (41, 42) or ribosome (43). In contrast, ASOs are becoming the method of choice to knock down lncRNAs, since they are thought to act on nascent RNA in chromatin (44). Given that cis-lncRNAs presumably act locally in chromatin, then one would expect ASO perturbations to have greater power to discover cis*-*lncRNAs. To test this, we evaluated predictions from each perturbation technology separately (Figure 3D). We observed broadly similar rates of cis*-*lncRNA identification between perturbation methods, supporting the notion that RNAi is readily active in the nucleus (45–47). However, ASO experiments appear to discover similar rates of activators and repressors, while RNAi perturbations yield an apparent excess of activators over repressors, together suggesting that differences do exist between RNAi and ASO perturbations.

While the small numbers preclude statistical confidence, these findings broadly support the use of RNAi in targeting nuclear lncRNAs and identifying cis-lncRNAs, although the possibility for perturbation-specific biases should be further investigated.

### Clues to cis-lncRNA mechanisms from localisation, expression and evolutionary conservation

We next sought insights into cis-lncRNA mechanisms by examining a range of features related to functionality and subcellular localisation. Although it has been previously postulated that cis-acting lncRNAs are more localized in the nucleus than the cytosol and have an overall lower RNA expression level (14), we found no evidence of any difference in nuclear/cytoplasmic localization between cis-lncRNAs and other lncRNAs (Figure 4A, Supplementary Figure S4A). Thus, cis-activity does not impact a lncRNA’s rate of nuclear export. Since many lncRNAs are non-polyadenylated, we similarly analysed nuclear/cytoplasmic localisation of total RNA (polyA+ and polyA-) in HepG2 and K562 cells. Here, we also observed no difference in localisation of cis-lncRNAs (Supplementary Figure S4C).

**Figure 4:**
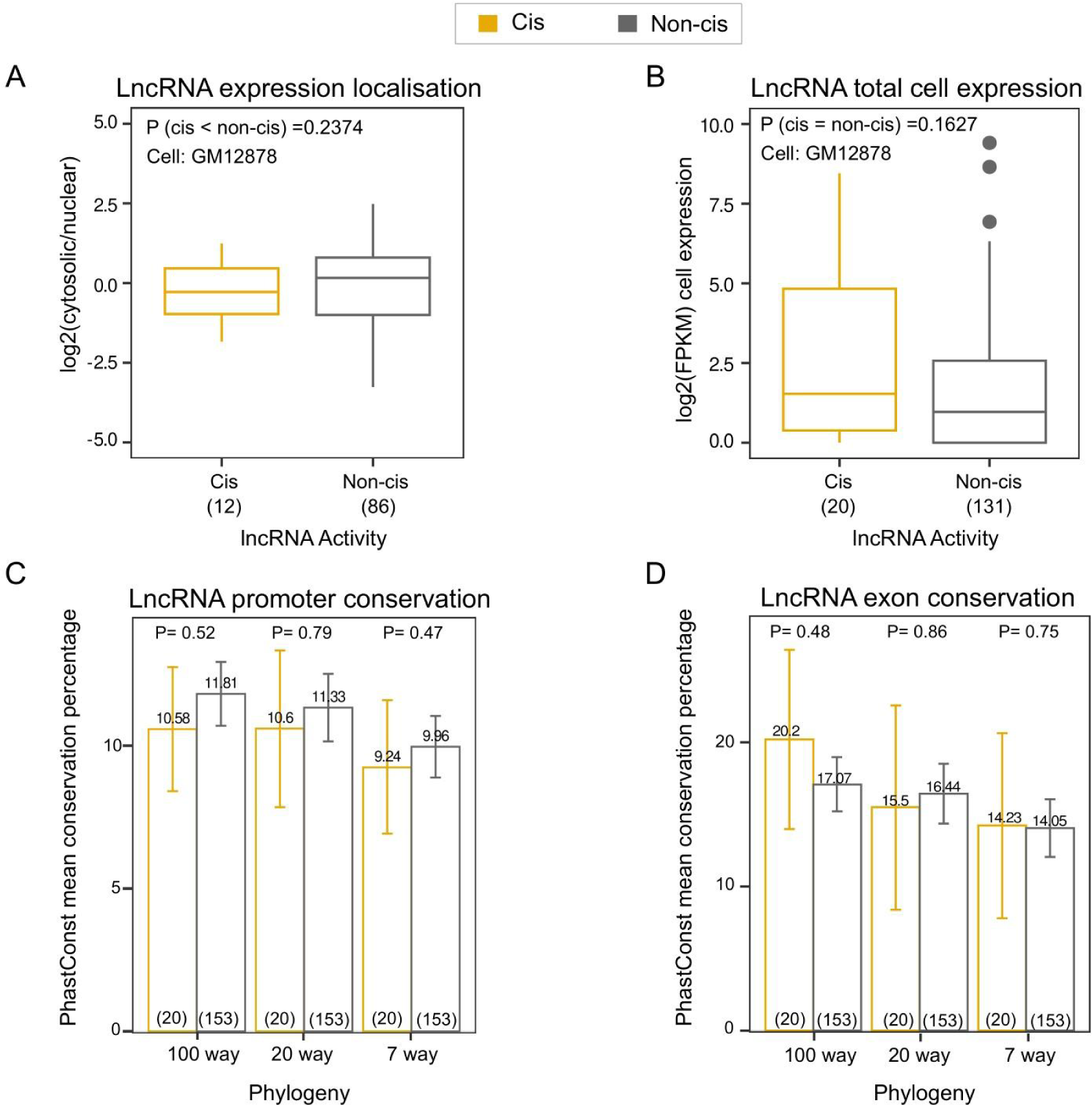
Subcellular localisation, expression and conservation of cis- and non-cis-lnRNAs. A) The distribution of the (log2-scaled) ratio of cytosolic to nuclear expression levels for cis- and non-cis-lnRNAs in GM12878 cells. The number of lncRNA genes in each group are displayed below, and represent the subset of lncRNA genes for which localisation data was available. Reported p-values for significance of between group differences are based on one-tailed Wilcoxon-rank-sum test. B) As for A) exploring the difference in levels of whole cell expression. Reported p-values for significance are based on two-tailed Wilcoxon-rank-sum test. C) and D) Barplots display the mean evolutionary conservation for the indicated features (x-axis). Number of genes in each group are displayed in brackets, and represent the subset of lncRNA genes for which PhastCons score was available for its promoter and exonic regions. Error bars represent standard deviation. Evolutionary conservation is calculated using PhastCons scores, where promoter conservation represents the percentage of promoter nucleotides overlapped by PhastCons conserved elements, and exon conservation represents the percentage of merged exonic nucleotides overlapped by PhastCons conserved elements. The reported p-values indicate the significance difference between cis- and non-cis-lnRNAs for each feature, determined using a one-sided Welch test with the alternative hypothesis of ‘greater’ (cis > trans).

Similarly, we evaluated the whole-cell expression level of cis-lncRNAs and observed a trend for average expression levels to exceed those of non-cis-lnRNAs in a number of cases, although these differences did not reach statistical significance (Figure 4B, Supplementary Figure S4B).

Evolutionary conservation and gene expression patterns are considered to yield important clues to lncRNA functionality (48). Promoter conservation and high expression are taken as general evidence for functionality (49), whereas conservation specifically in exons is reflective of the functionality of mature RNA transcripts (50). Some mechanistic models for cis-lncRNAs posit that they act through non-sequence dependent features (14), predicting that cis-lncRNAs’ exons display background levels of evolutionary conservation, whereas their promoter regions (controlling expression) are more conserved. To test this, we evaluated the evolutionary conservation for promoter and exons for three different vertebrate phylogenies (Figure 4C-D). This revealed no discernible difference in exonic or promoter conservation for cis-lncRNAs over non-cis-lnRNAs, and suggesting that cis-lncRNAs’ nucleotide sequence is important for the functionality. A lack of statistical power due to a low sample size and high variance should be noted.

Finally, we asked whether proximal gene targets of cis-lncRNAs are more strongly regulated, compared to distal gene targets. We compared the fold change in gene expression of proximal and distal targets for each cis-lncRNA (Supplementary Figure S5A). This revealed that distal and proximal targets tend to have similar degrees of regulation, however with notable exceptions including LINC01615 (activator) and DNAAF3 (repressor), where proximal targets are significantly more strongly regulated.

Together, these findings suggest that cis-lncRNAs are broadly similar to other lncRNAs in terms of expression, subcellular localisation and overall functionality.

### Association of cis-lncRNAs with enhancer elements

It has been widely speculated that cis*-*lncRNAs, particularly activators (ie e-lncRNAs), act in concert with DNA enhancer elements to upregulate target gene expression (3, 9, 12). Our catalogue of cis-lncRNAs represents an opportunity to independently test this. To do so, we calculated the rate of overlap of lncRNAs with enhancers using epigenomics data across human tissues (Figure 5A-B, Supplementary Figure S6). Analyses were performed at a variety of epigenome thresholds (the minimum number of samples required to define a given epigenomic state) and window sizes (the distance from the lncRNA TSS to the nearest epigenome element).

**Figure 5:**
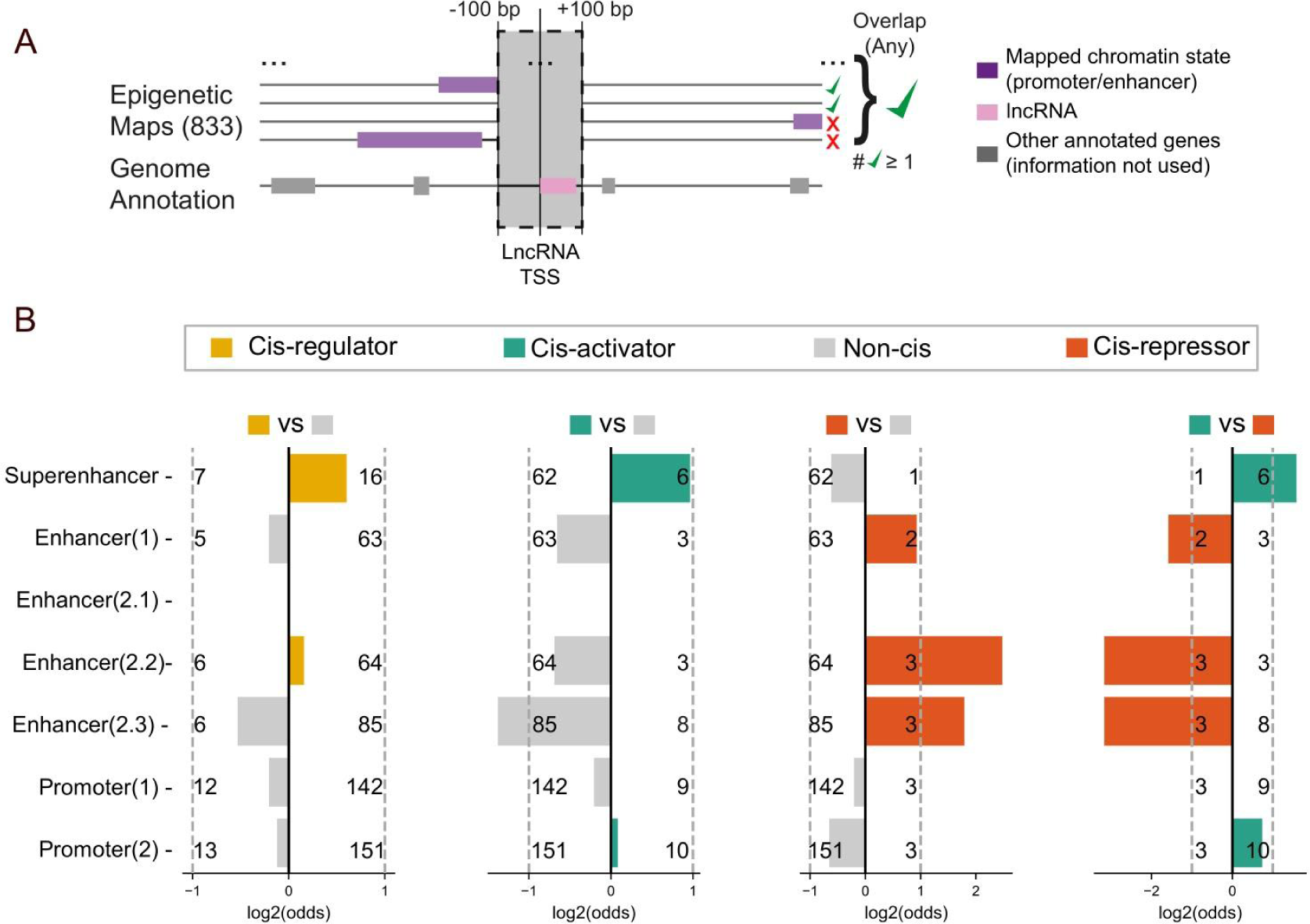
Overlap of cis-lncRNAs with enhancer elements. A) Method of calculating overlap by enhancer annotations (horizontal purple bars) of lncRNA TSS (pink bar). Overlaps are considered while varying two thresholds for defining a lncRNA-enhancer overlap: minimum numbers of individual enhancer annotations (epigenome threshold) and window size. Only the TSS spans with overlaps in more samples than a given epigenome threshold are considered. B) Enrichment results for epigenome threshold=1 and span=100 bp. Rows show enrichment for super-enhancer, enhancer, and promoter states while comparing the TSS according to their mechanism of action (see Methods).

This analysis revealed several intriguing trends of association between cis-lncRNAs and enhancer elements, although none reached statistical significance at the given sample size (Figure 5B). Broadly, we observed a generalised enrichment of various enhancer element annotations with cis-lncRNAs, notably the cis-activators with Superenhancers, and the cis-repressor with Enhancer(2.2) and (2.3) elements. Inspection of overlaps at other thresholds and window sizes revealed a similar effect (Supplementary Figure S6). Within the limits of statistical power given our relatively small sample size, these findings suggest only a weak relationship between cis-lncRNAs and enhancer elements that should be re-examined in future studies.

### Some cis-lncRNAs are brought into spatial proximity to their targets by chromatin looping

A second mechanistic model posits that regulatory interactions between cis-lncRNAs and target genes are defined by spatial proximity, brought about by chromatin looping (Figure 6A). Once again, our cis-lncRNA catalogue makes it possible to test this. To measure proximity, we utilised published Hi-C interactions from a range of human cell lines (30). We evaluated the importance of proximity for regulatory targeting, by combining an asymptotic regression model to predict an “expected interaction” at a given linear genomic distance, with a logistic regression model to evaluate whether strong deviations from this expectation were indicative of targeting (Figure 6). This approach revealed a significant (-log10(P)≥ 1.3) contribution of spatial proximity to targeting for cis-activator lncRNA UMLILO (8 cell lines) (Figure 6). An alternative approach (inverse square model) confirmed this result and additionally yielded DA125942 (8 cell lines) (Supplementary Figure S7A,B). Further confidence in these results comes from the fact that, for both lncRNAs, previous studies have implicated chromatin looping in their targeting mechanism (8, 37). An excellent example is represented by HUVEC cells, where UMLILO target genes tend to be located in higher proximity (Interaction, y-axis), compared to other non-targets at similar distances in linear DNA (x-axis) (Figure 6C, Supplementary Figure S7C). Together, this indicates that for a subset of cis-lncRNAs, spatial proximity brought about by chromatin looping may determine the identity of target genes.

**Figure 6:**
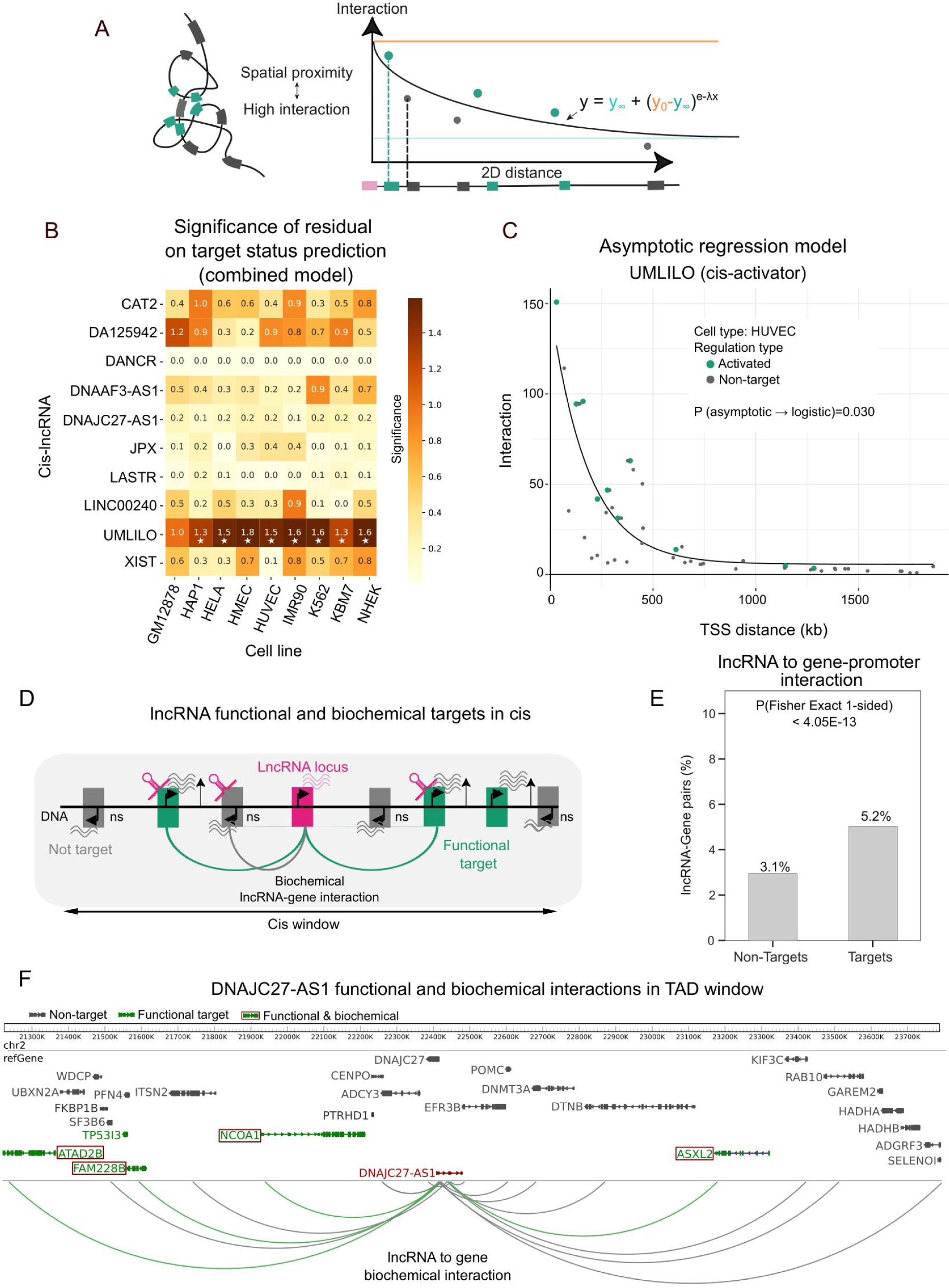
Linking lncRNAs to target genes with DNA:DNA chromatin looping RNA:DNA and physical association. A) A model for proximity-driven target selection: (Left panel) Chromatin folding brings lncRNA (pink) into spatial proximity with proximal genes, which are subsequently targeted (green). (Right panel) Chromatin proximity maps, such as provided by HiC methodology, enable one to evaluate the spatial proximity (y-axis) of targets, while normalising for confounder of linear 2D DNA distance (x-axis). These parameters were modelled using an Asymptotic regression model (right panel, inset). B) Evaluating the contribution of proximity to target selection in human cells: The model significance of cis-lncRNAs (identified by TransCistor-digital) (x-axis) was evaluated across HiC interaction data from a panel of human cell lines (y-axis). Colour scale shows uncorrected p-values; Asterisks represent P<0.05. Darker colors cells indicate cases where target genes tend to be significantly more proximal than non-targets. No cases of the inverse were observed. C) Example data for UMLILO in HUVEC cells. Note that target genes (green) tend to be more spatially proximal (y-axis) than non-target genes (grey) at a similar TSS-to-TSS genomic distance (x-axis). D) A hypothesis for biochemical interaction of lncRNAs with their functional target genes: lncRNA transcripts (pink) are transcribed from their gene locus and physically recruited to target genes (green). Non-specific or non-productive recruitment is also observed at non-target genes (grey), yet at a lower rate. E) Rate of observed biochemical interactions between lncRNAs and same-chromosome genes (y-axis), comparing lncRNA-target relationships classified as functional ‘target’ and ‘non-target’ from our data. Biochemical interactions were collected from published experimental measurements (see Materials and Methods). Reported p-values for significance are based on one-sided Fisher’s exact test. F) Example data for DNAJC27-AS1 (red) depicting observed biochemical interactions with nearby functional target genes (in green) and non-target genes (in grey) within its TAD domain. The target genes linked to the lncRNA by biochemical interaction are marked with a red box.

### Cis-lncRNA transcripts are biochemically bound to their target genes

Recent studies have mapped RNA:DNA contacts at a global scale and have demonstrated that the majority of contacts occur locally to the RNA transcription, suggesting a mechanism for cis-regulation of genes by ‘biochemical’ contact of a lncRNA transcript (51–53). To examine this further, we evaluated the overlap of functional lncRNA-target relationships (defined by LOF experiments) and biochemical relationships (defined by RNA:DNA contacts). Using the entire human lncRNA set (cis- and non-cis-), we assessed significant lncRNA-to-DNA contacts forall genes promoters on the same chromosome (Figure 6D). This revealed that functional target genes are significantly more likely to also be biochemically bound by their regulator lncRNA, compared to non-target genes (P<4.05E-13) (Figure 6E).

An illustrative case is DNAJC27-AS1 (Figure 6F), identified as a cis-activator by TransCistor-digital. Within its local TAD, DNAJC27-AS1 has significantly more biochemical interactions with its functional target genes (4 out of 5) compared to its non-targets (11 out of 30) (P=0.04, Fisher’s exact test, 1-sided). Overall, these findings suggest that functional regulation of target genes is determined, at least in part, by biochemical recruitment of the regulatory lncRNA.

## DISCUSSION

We have described TransCistor, a pair of quantitative methods for the identification of cis-regulatory lncRNAs. We applied it to a corpus of perturbation datasets to create the first large-scale survey of cis-regulatory RNAs. We evaluated the performance of TransCistor in light of the present state-of-the-art and used the resulting catalogue of cis-lncRNAs to address fundamental questions regarding their prevalence and molecular mechanisms.

TransCistor-digital and -analogue represent practical tools for cis-lncRNA discovery. TransCistor enables researchers to identify cis-regulatory lncRNAs, based on the distribution of target genes identified through a perturbation and transcriptomic analysis. The definition of cis-lncRNAs that we have here formulated is rigorous yet also consistent with the field, which has since the discovery of *XIST* and *H19* employed a functional definition of cis-regulation: “*Cis*-acting lncRNAs… regulate gene expression in a manner dependent on the location of their own sites of transcription”(14).

Their two distinct statistical methods are designed to capture a range of cis-activity, from lncRNAs regulating the most proximal neighbour gene’s expression within the local TAD, such as *CHASERR* (*4*), to those regulating a more distal target amongst other non-target genes, such as *CCAT1-L* (7). The value of resulting predictions is supported by good statistical behaviour as judged by quantile-quantile (QQ) analysis, consistency between methods and datasets, and recall of numerous known cis-lncRNAs, including founding members *H19* and *XIST*. TransCistor is made available both as a webserver and standalone software. It is compatible with a wide range of input data, since “regulation” files can be readily generated from any experimental dataset comprising lncRNA perturbation and global readout of gene expression changes, including two decades of experiments from microarrays to RNA-sequencing and cap analysis of gene expression (CAGE). Importantly, TransCistor is ready to deploy with recently-developed and future parallelised CRISPR LOF methods such as Perturb-Seq (54), raising the possibility of comprehensive mapping of cis- and trans-regulatory lncRNAs. It is important to note, however, that identification of cis-target genes requires whole transcriptome datasets, meaning that signature methods based on subsets of genes, such as LINCS (55), will not be suitable.

This work builds on important previous attempts to comprehensively discover cis-regulatory lncRNAs. Basu and Larsson utilised gene expression correlation as a means for inferring candidate cis-regulatory relationships (56). Very recently, de Hoon and colleagues employed genome-wide RNA-chromatin and chromatin folding to train a predictive model for cis-regulatory lncRNAs (57). While these methods are valuable, they infer target genes based on indirect correlates of cis-regulation, which often do not reflect causation (58). Furthermore, we could only find evidence that chromatin folding links cis-lncRNAs to their target genes in a minority of cases. What distinguishes TransCistor from these approaches, is its use of LOF perturbations to directly identify gene targets. We argue that, due to its direct and functional nature, this approach should be considered the gold standard evidence for defining cis-regulatory relationships. Moreover, TransCistor’s versatile nature allows its widespread adaptation which, in the future, could be used to build machine learning models of cis-action utilizing numerous features, such as expression, fold change of targets and distance.

A key insight from this work is the low statistical power available to identify cis-lncRNAs and the consequent high rate of false-positive predictions. Previous studies used a “naïve” criterion of ≥1 cis-target gene within an arbitrarily-sized window; however, we show that this method is prone to predominantly false-positive predictions at ≥50 kb windows. TransCistor improves on this situation by making predictions at a defined false discovery rate (FDR). Nevertheless, the statistics of cis-lncRNA prediction depend on the distribution of target and non-target genes around the lncRNA in question. This means that statistical power to identify cis-lncRNAs is inherently constrained by biology. Several likely examples of false positive predictions are well-studied lncRNAs *NORAD* and *DANCR*, which are represented by numerous perturbation experiments in our dataset, where only one was called a hit. Conversely, several factors likely contribute to false negatives, including low statistical sensitivity (i.e. presence of few cis-targets relative to trans-targets) and correction for multiple testing. Furthermore, it is likely that direct target mRNAs in turn regulate numerous, indirect downstream genes, creating a high background that further reduces detection sensitivity. This latter issue might be addressed in future by performing transcriptomic analysis at short timepoints immediately after lncRNA LOF to observe direct target genes. Finally, the majority of lncRNAs were tested in only a single cell type, and it is entirely possible that lncRNAs display cis-regulatory activity in a cell-type specific manner. These considerations should prove useful in the design of future experiments to identify cis-lncRNAs. To minimise false-positive predictions, we recommend that colleagues perform multiple independent perturbations (e.g. two or more distinct ASO sequences), perform transcriptomic analysis at earliest possible timepoints, and only consider cis-lncRNAs on the basis of ≥2 consistent results.

Our results afford important insights into the regulatory lncRNA landscape. Notwithstanding the caveats discussed above, we provide the first global estimate of cis-lncRNA prevalence, suggesting they represent a modest fraction (8%) of the total, with a slight prevalence of activators over repressors. These values are certainly impacted by a variety of errors discussed above, which we hope will be corrected by future, larger-scale studies. The preponderance of cis-activators may be an artefact of RNAi perturbations, which appear to yield an excess of activators over repressors. Our results shed light on cis-lncRNAs’ molecular mechanisms, finding evidence challenging the notion that spatial proximity defines lncRNA targets. Surprisingly, cis-lncRNAs are not preferably localized to the nucleus, nor are more evolutionarily conserved or more expressed than non-cis-acting lncRNAs. On the other hand, we find non-significant associations of cis-lncRNAs with enhancer elements, which may point to a mechanistic relationship that awaits further investigation with greater sample sizes. Most important, perhaps, is the finding that lncRNA target genes are more likely to have evidence for physical association of the lncRNA transcript. This implies that, at least some lncRNAs regulate downstream mRNAs by physically associating with their gene. Overall, these findings show the utility of cis-lncRNA catalogues in examining molecular mechanisms of regulation, although larger datasets will be required in future to draw more conclusive inferences than could be done here.

Finally, it is worth revisiting the assumptions we make when interpreting lncRNA perturbation experiments. These involve a small oligonucleotide with perfect sequence complementarity to a lncRNA target in both RNA *and* DNA, and assess the outcome in terms of steady-state RNA levels. Two key assumptions are made. Firstly, any change in downstream gene expression is assumed to occur through changes in the targeted lncRNA transcript. It is well known that small oligos are not only capable of hybridising to genomic DNA (59) but also affecting local chromatin modifications (60), raising the possibility of chromatin/DNA-mediated cis-regulatory mechanisms that could be misinterpreted as lncRNA-mediated effects. The second assumption is more fundamental:, when local gene changes are observed to occur, such changes reflect the biological function of the lncRNA (61, 62). The alternative explanation is that perturbations of a lncRNA lead to changes to local gene expression, but that this is a by-product of altering lncRNA expression (e.g. by disrupting local transcription factories), and that the evolutionarily-selected function of the lncRNA is something quite different. In other words, is observed cis-activity a reflection of a genuine, adaptive biological regulatory pathway, or is it merely a technical artefact without biological relevance? Testing these alternative explanations will be an interesting challenge for the future, facilitated by the tools provided here.

## DATA AVAILABILITY

Most of the data underlying this article were derived from sources in the public domain: FANTOM (https://fantom6-collaboration.gsc.riken.jp//webdav/home/), lncRNA2Target (http://bio-annotation.cn/lncrna2target/). XIST knockdown RNA-sequencing data were kindly provided by Hyeonjoo Lee and Jin-Wu Nam (Hanyang University, Republic of Korea). TransCistor source code, all regulation files and related information can be found on GitHub (https://github.com/gold-lab/TransCistor). The webserver can be accessed at https://transcistor.unibe.ch/.

## SUPPLEMENTARY DATA

Supplementary Data (Supplementary File 1) are available at NAR online.

## ACKNOWLEDGEMENTS

We thank the other members of GOLD Lab for insightful feedback and discussions. Hyeonjoo Lee and Jin-Wu Nam (Hanyang University, Republic of Korea) kindly shared Xist knockdown RNA-sequencing data. All computation was carried out on the University of Bern Interfaculty Bioinformatics Unit cluster maintained by Rémy Bruggman and Pierre Berthier. We acknowledge administrative and logistical support from Basak Ginsbourger, Rahel Tschudi, Marla Rittiner, Beatrice Stalder, Willy Hofstetter and Patrick Furer (DBMR, University of Bern).

---

**Supplementary Figure S1.**
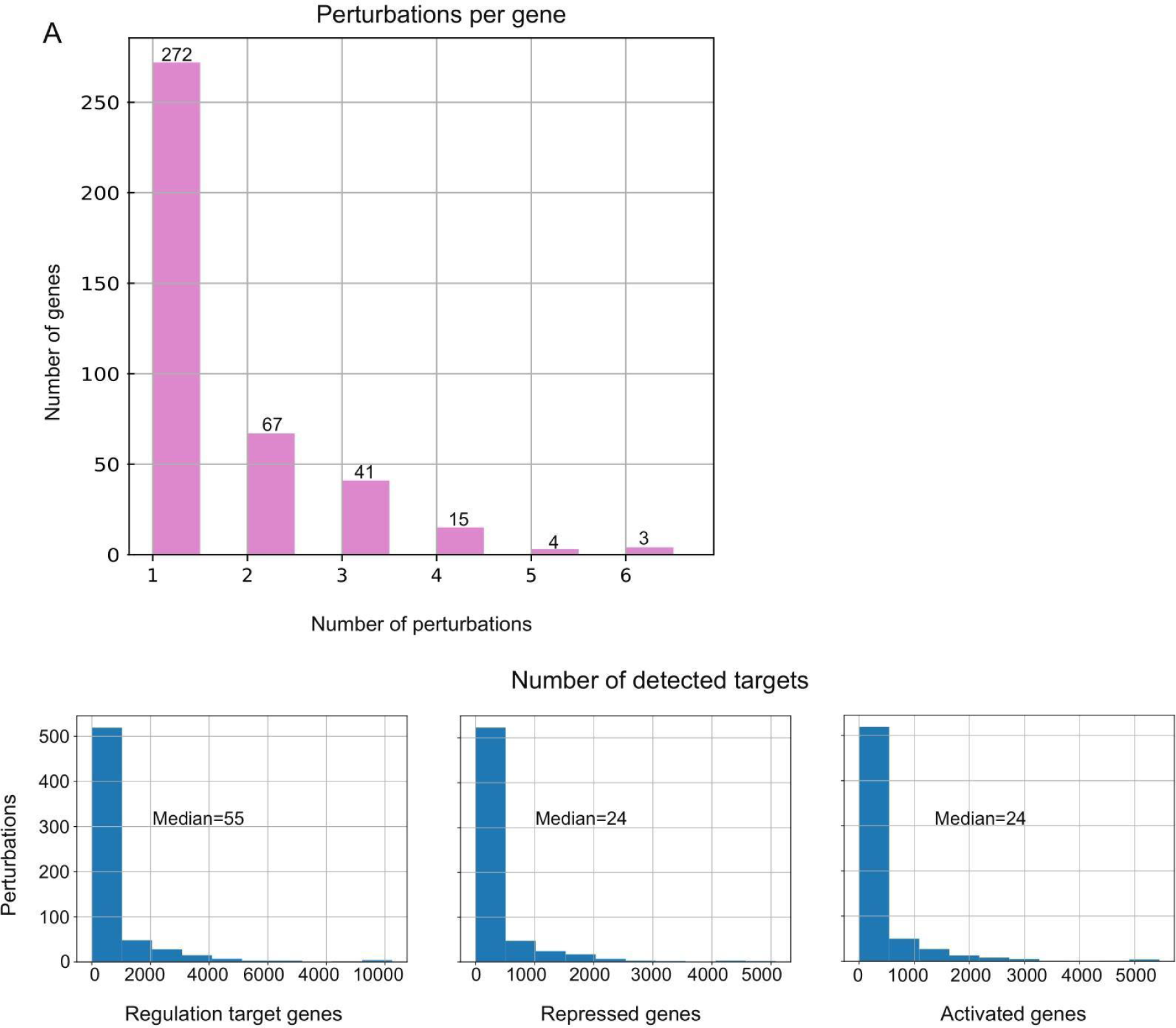
Summary statistics of perturbation datasets. A) Histogram displaying the numbers of separate perturbation experiments (x-axis) available for each lncRNA gene (y-axis). B) Histograms displaying the number of significantly changing genes (targets) (x-axis) for each perturbation experiment (y-axis). Regulated genes represent the union of activated and repressed genes.

**Supplementary Figure S2.**
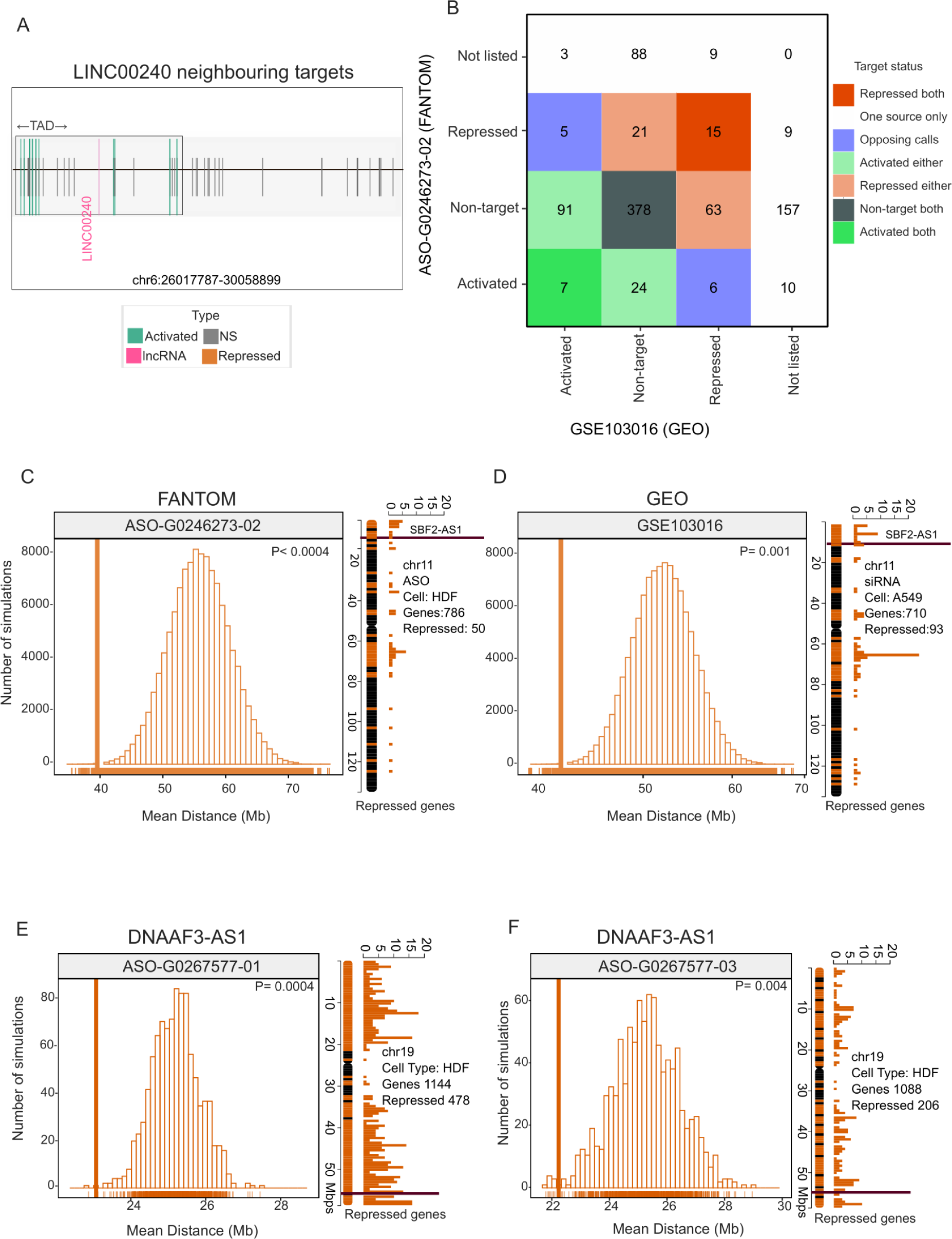
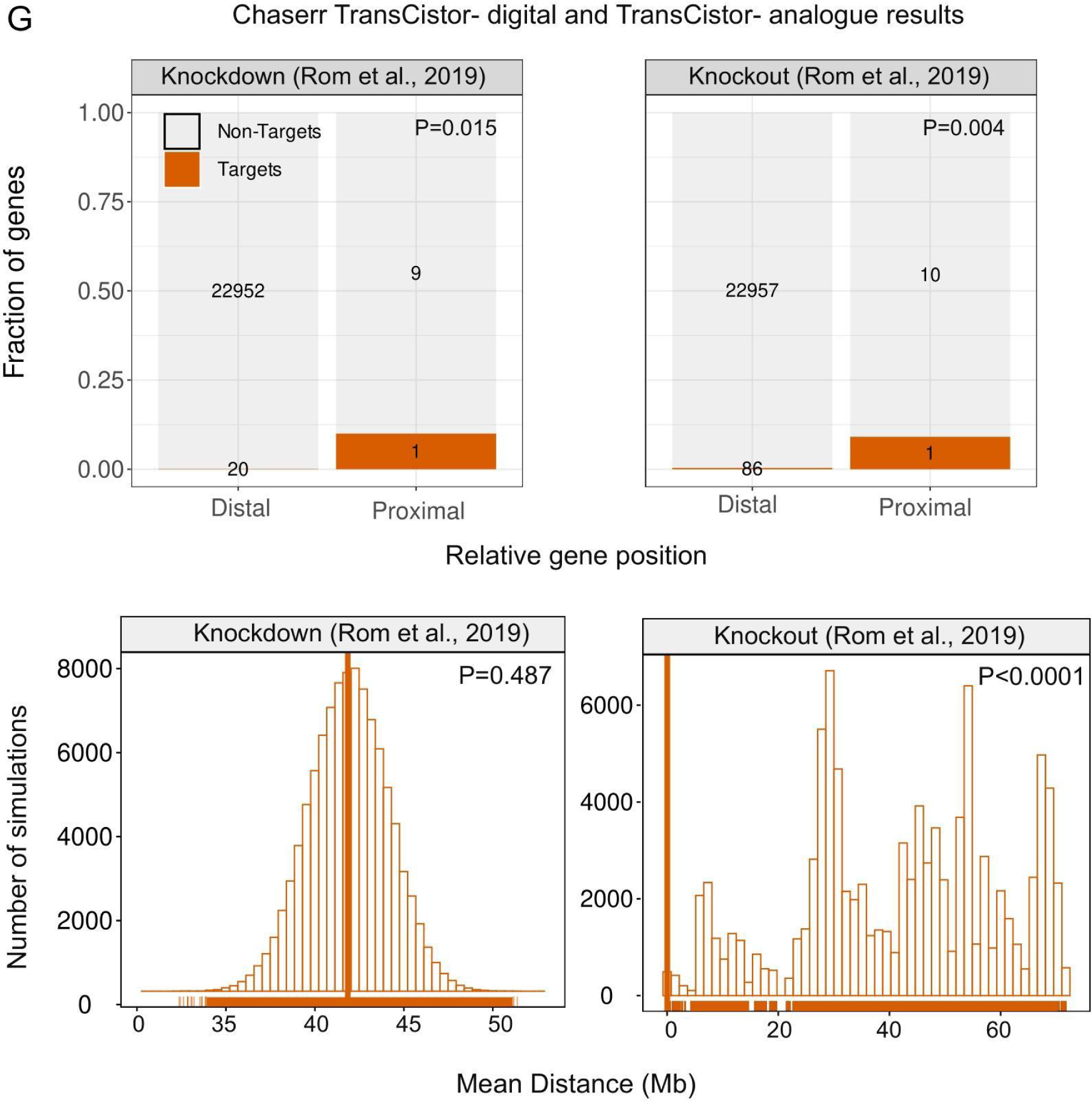
Exploring independent and supporting datasets for significant cis lncRNAs. A) LINC00240 genomic locus: Vertical bars denote gene TSS. Grey: non-targets; green: activated targets; pink: LINC00240. Black box: Topologically associated domain. B) SBF2-AS1 target regulation overlap between FANTOM ASO knockdown and independent siRNA GEO dataset. Numbers indicate the genes in each category, classified by their regulation in the two distinct datasets. C) TransCistor-analogue results for FANTOM ASO knockdown targeting SBF2-AS1 in human dermal fibroblasts. D) As for (C), but for independent data from A549 cells treated with siRNA. E) DNAAF3-AS1, an example cis-repressor identified by TransCistor-analogue. Shown are the target distance statistic (x-axis) for real data (vertical bar) and simulations (boxes). The number of simulations in each distance bin is displayed on the y-axis. F) As for (E), for a second perturbation experiment. G) Chasser lncRNA comparison with TransCistor-Digital (top) TransCistor-Analogue (bottom) across multiple datasets and methods; TransCistor-Digital (top) plot shows the number of genes, divided by targets / non-targets (colour / grey), location (distal/proximal) and regulation direction (activated/repressed). TransCistor-Analogue (bottom) plot show target distance statistic (x-axis) for real data (vertical bar) and simulations (boxes). The number of simulations in each distance bin is displayed on the y-axis. Chasser is classified as a cis-repressor, due to the significant excess

**Supplementary Figure S3:**
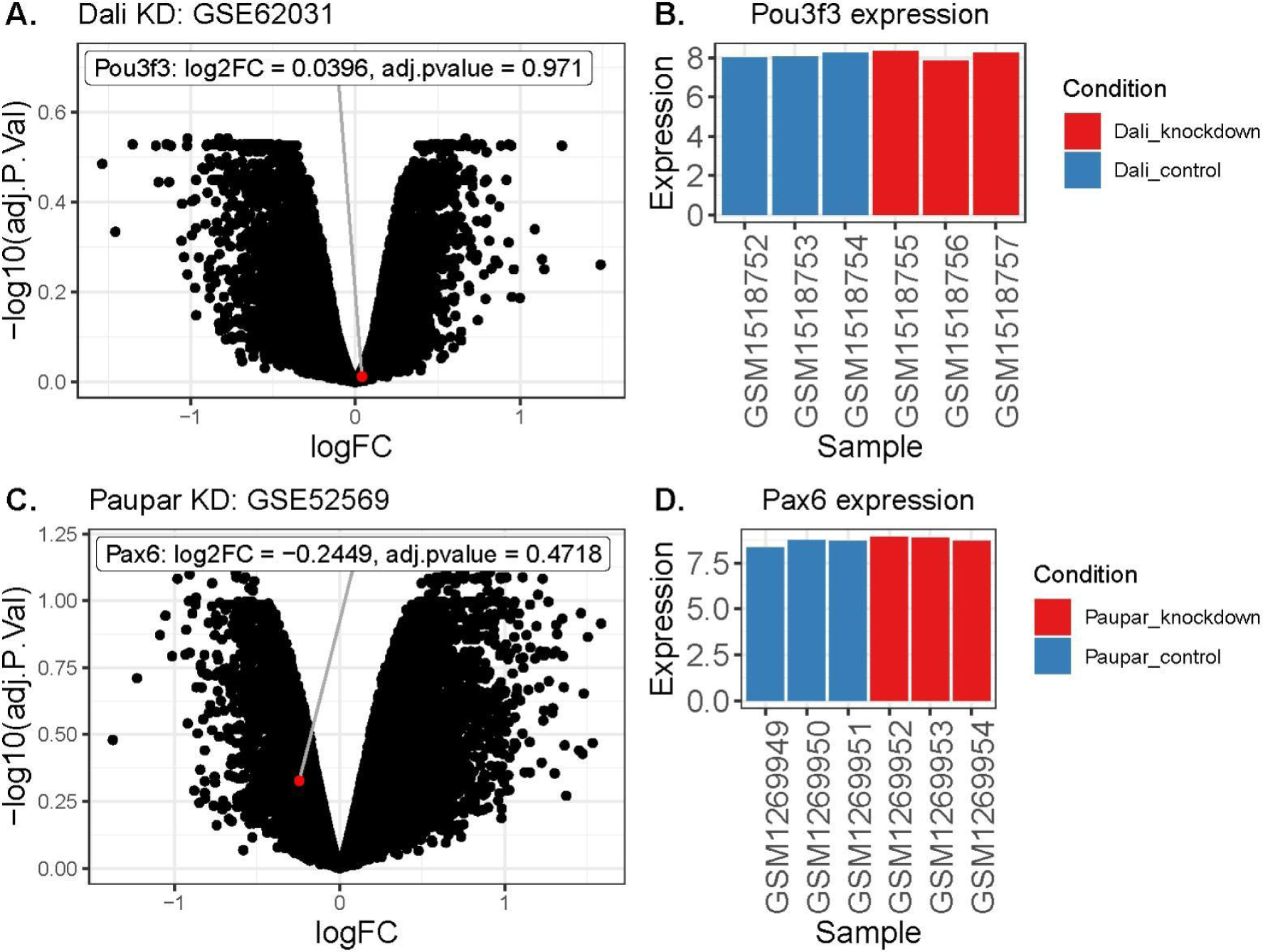
Analysis of Dali and Paupar target genes using public microarray data. A) Global transcriptome changes upon Dali knockdown were obtained from Gene Expression Omnibus (https://www.ncbi.nlm.nih.gov/geo/query/acc.cgi?acc=GSE62031). B) Neighbour gene and putative target Pou3f3 mRNA expression in control and Dali knockdown samples. C) Global transcriptome changes upon Paupar knockdown were obtained from Gene Expression Omnibus (https://www.ncbi.nlm.nih.gov/geo/query/acc.cgi?acc=GSE52569). D) Neighbour gene and putative target Pax6 mRNA expression in control and Paupar knockdown samples.

**Supplementary Figure S4:**
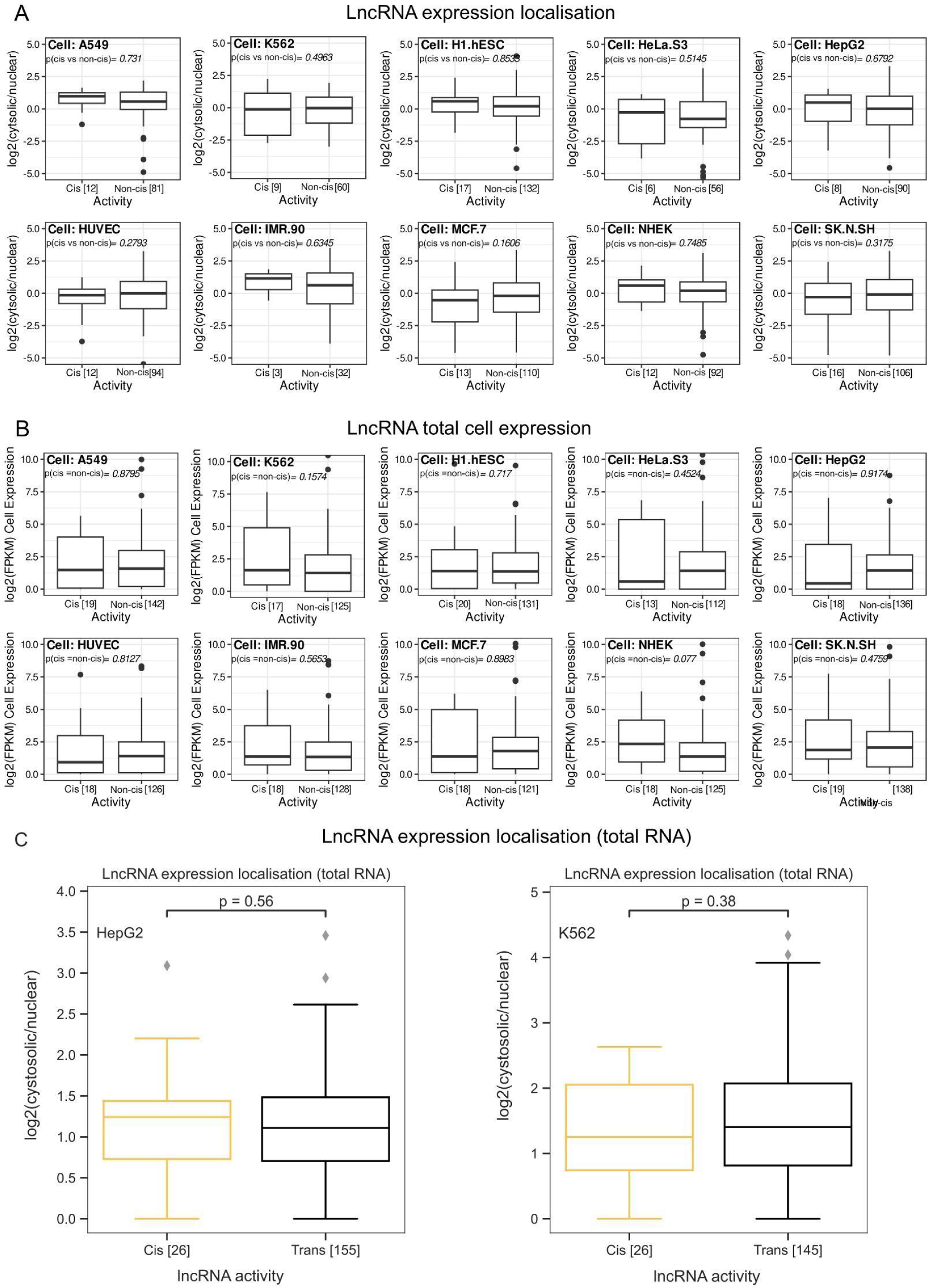
Cytosolic/nuclear localisation of cis/non-cis lncRNAs. Boxplots showing the distribution of the (log2-scaled) ratio of cytosolic to nuclear expression levels for cis-/ vs. non-cis-regulatory lncRNAs in different cell types. The number of genes in each group is displayed below and represents the subset of lncRNA genes for which localisation data was available. Reported p-values for the significance of between-group differences are based on a one-tailed Wilcoxon-rank-sum test (alternative: cis > non-cis). B) As for A) exploring the difference in total levels of cell expression. Reported p-values for significance are based on a two-tailed Wilcoxon-rank-sum test. C) The distribution of the (log2-scaled) ratio of cytosolic to nuclear expression levels for cis- and non-cis-lnRNAs in HepG2 (left) and K562 (right) cells. Number of genes in each group are displayed below, and represent the subset of lncRNA genes for which localisation data was available. Reported p-values for significance of between group differences are based on one-tailed Wilcoxon-rank-sum test.

**Supplementary Figure S5:**
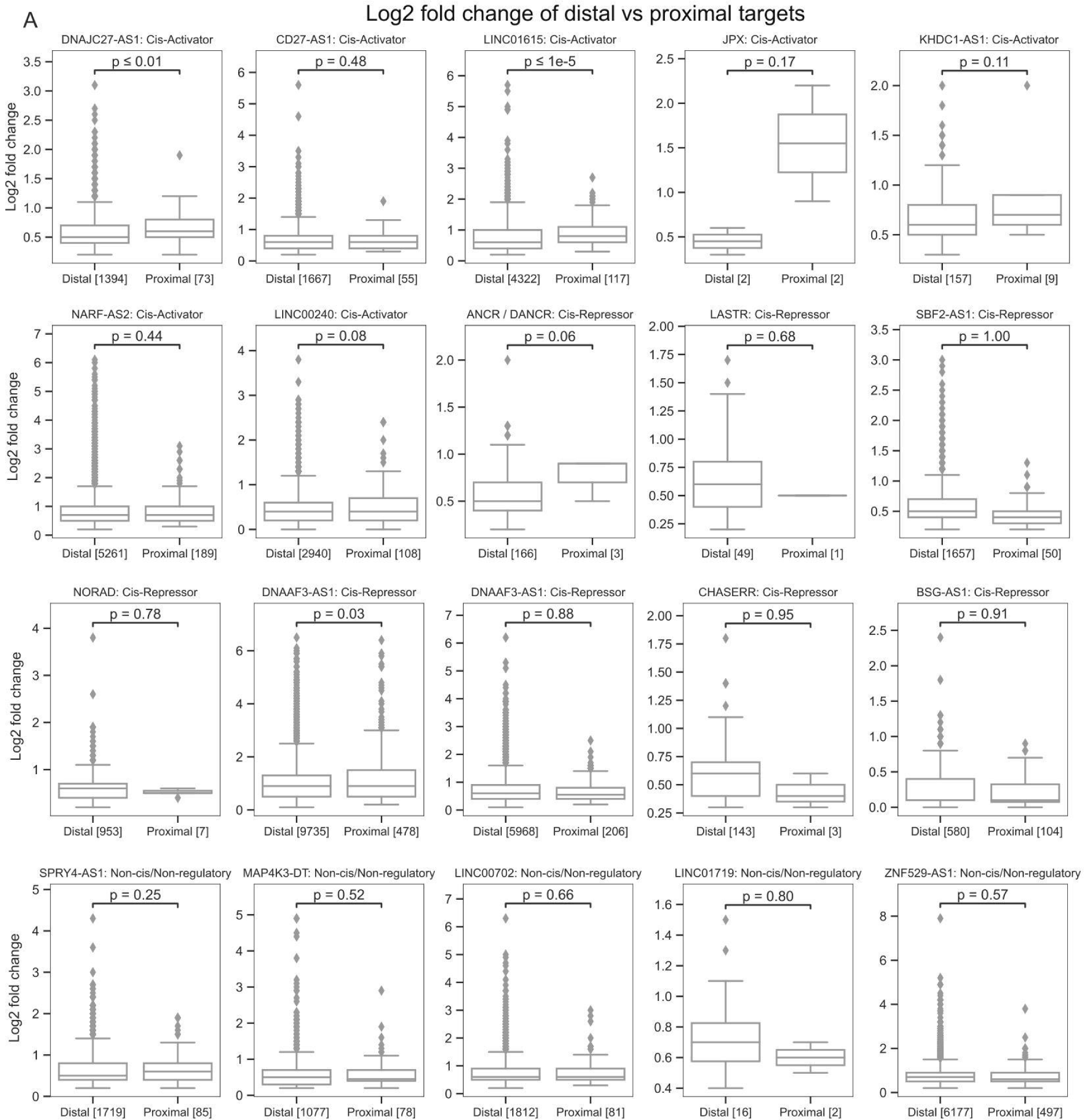
Foldchange difference between proximal and distal gene targets. A) Exploring log foldchange difference of lncRNA targets in proximal vs distal for cis-activators, cis-repressors and randomly selected non-cis lncRNAs. Reported p-values for significance of between group differences are based on one-tailed Mann–Whitney test.

**Supplementary Figure S6:**
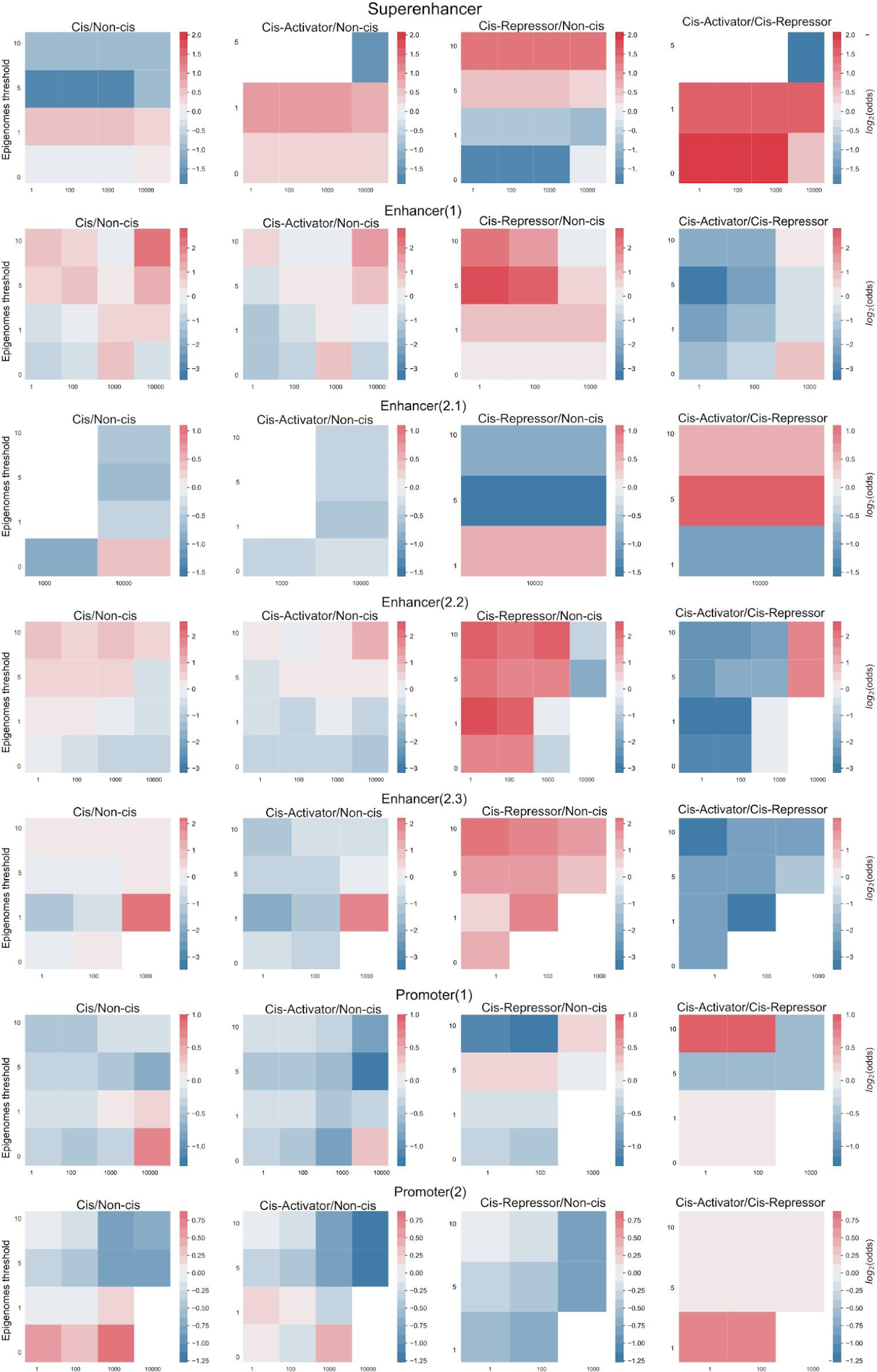
Enrichment of enhancers, super-enhancers and promoters in *cis*-lncRNAs. Each row represents a different enhancer annotation (see Methods). Columns represent comparisons between indicated pairs of lncRNA classes. Heatmaps display the enrichment of overlap at different genomic windows around the lncRNA TSSs (span, *x*-axis) and the minimum number of observed samples required to define an enhancer (epigenome threshold, *y*-axis). Statistical significance is indicated where p-value ≤ 0.05 by Fisher’s exact test (1-sided), and not corrected for multiple hypothesis testing.

**Supplementary Figure S7:**
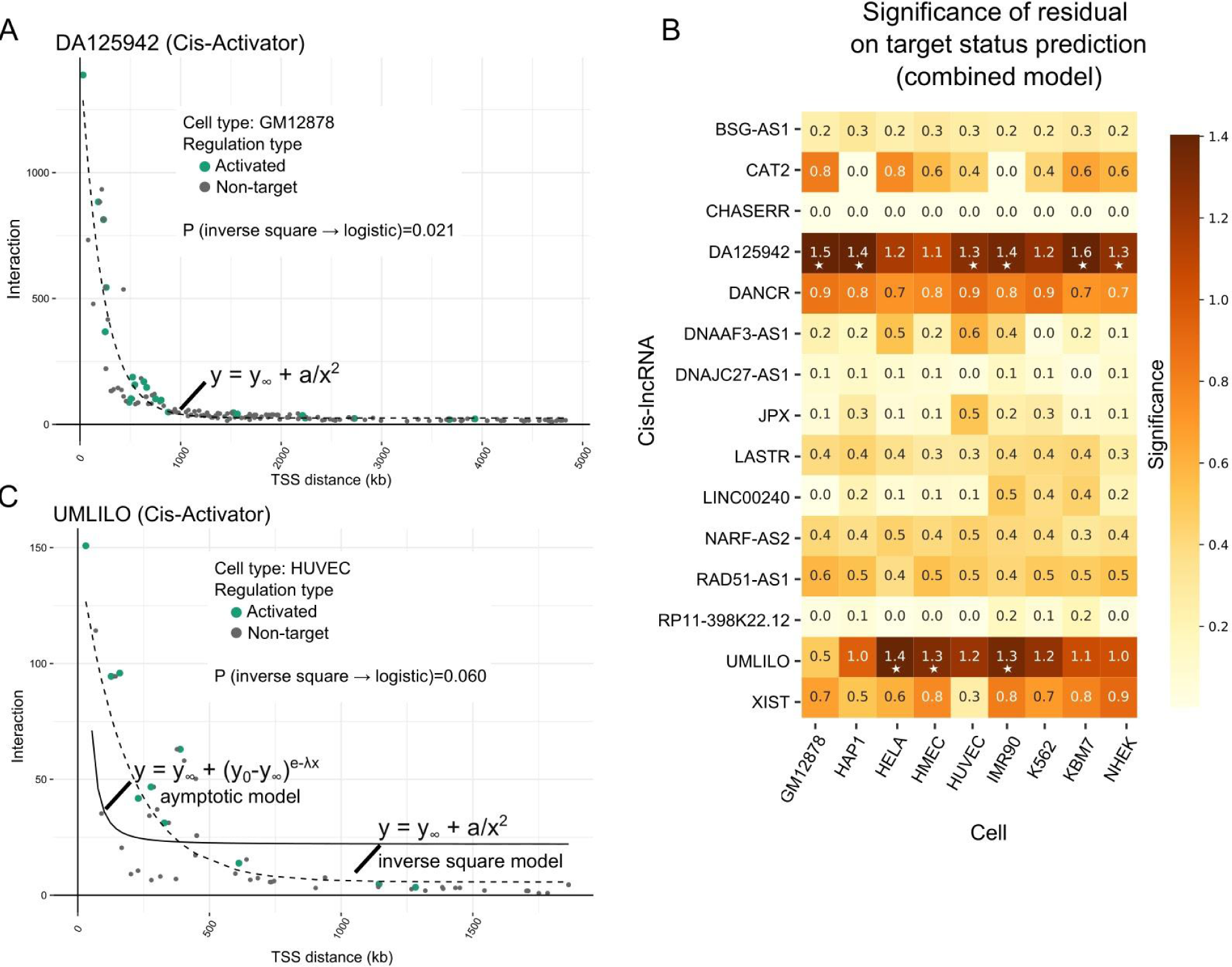
Alternative inverse square model for target gene interaction. A) and B) are equivalent to main Figure 5 panels E and D respectively but using an inverse square model instead of an asymptotic regression. C) Comparison of asymptotic (solid line) and inverse square (dashed line) regression models.

## Supplementary Files

**Table S1. Number of experiments and unique lncRNAs**

**Table S2. Individual experiment information and TransCistor predicted activity**

**Table S3. Existing literature on the cis-activity of the lncRNAs tested**

